# A Tether-Independent Mechanism for Bacterial Outer Membrane Separation

**DOI:** 10.1101/2025.05.15.654258

**Authors:** Carolina Basurto De Santiago, Carlos A. Ramírez Carbó, Govind Prasad Sah, Daniel Wall, Shiwei Zhu, Beiyan Nan

## Abstract

Gram-negative bacteria divide by coordinating the separation of two cell envelope layers: the peptidoglycan (PG) sacculus and the outer membrane (OM). In several well-studied model organisms, the OM is tightly tethered to the PG layer, causing both structures to constrict synchronously during cell division. In contrast, division of the PG and OM is uncoupled in *Myxococcus xanthus*, where the OM undergoes only limited invagination and remains largely unconstricted even after PG fission is complete, separating substantially later than the PG layer. Strengthening the OM–PG connection by overexpressing either the native tethering protein Pal or the heterologous lipoprotein Lpp restores synchronized constriction of the two envelope layers. These findings suggest that the strength of OM–PG coupling determines OM division dynamics and that the OM can divide by simple mechanical forces independent of OM–PG attachments.

## Introduction

The cell wall of Gram-negative bacteria consists of two distinct layers: a peptidoglycan (PG) meshwork and an outer membrane (OM). PG is a single macromolecule of glycan strands crosslinked by short peptides. The OM is an asymmetric bilayer with lipopolysaccharides (LPS) in the outer leaflet and phospholipids in the inner leaflet^1^. While PG plays a major role in maintaining cell integrity and morphology, the OM serves as a selective barrier that protects the cell from harmful substances, including antibiotics^2–4^. In some bacteria, such as *Escherichia coli*, OMs are load-bearing structures that can exhibit even greater stiffness than PG^5,6^. Recent studies reveal that OMs and PG function as an integrated cell envelope rather than independent layers^6^. Whereas PG synthesis has long been a primary target for antibiotic development, OMs are increasingly recognized as promising targets for antibacterial therapies and as platforms for vaccine development^1^.

During cell division, Gram-negative bacteria must separate both the PG and OM layers. This process can be conceptually divided into three stages: (i) constriction, where PG is assembled inward at division site, leading to localized narrowing of cell body; (ii) PG fission, marked by inner membrane (IM) separation; and (iii) OM fission (**Fig. 1**).

**Fig. 1.**
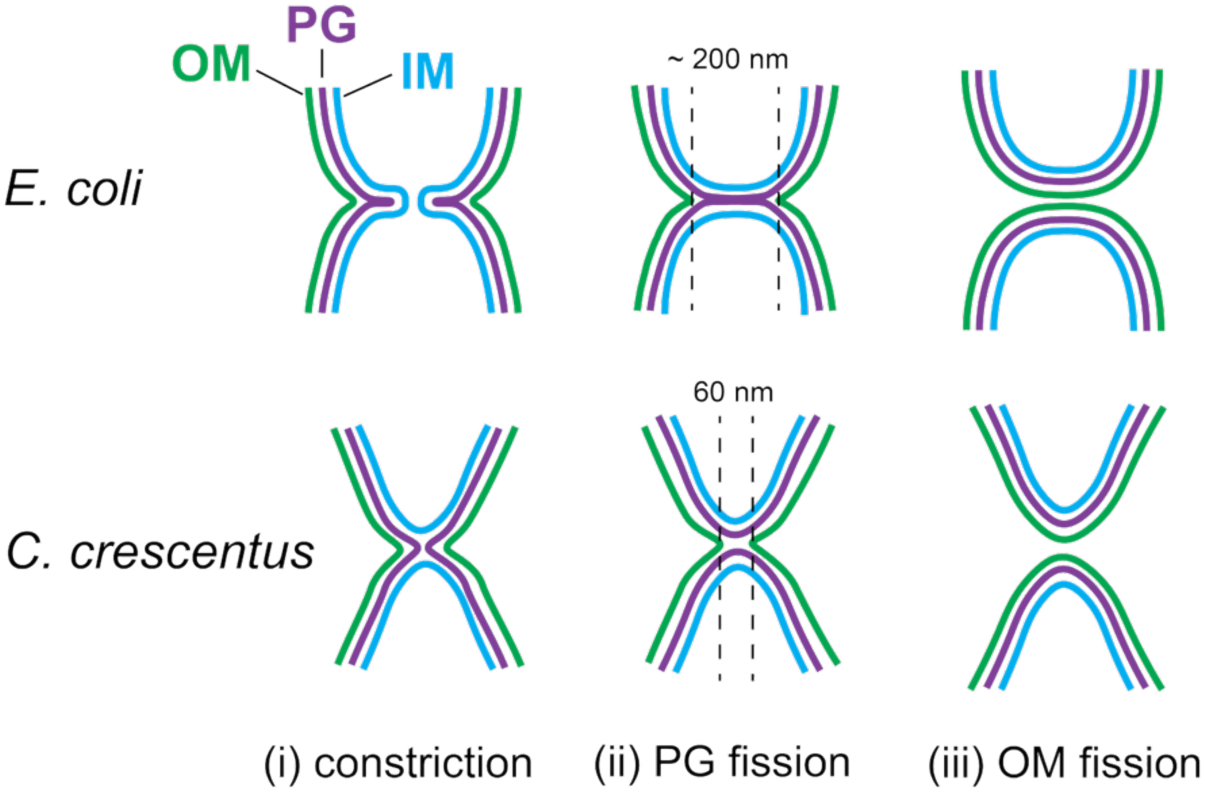
Variation in division mechanisms between *E. coli* and *C. crescentus*. Although *E. coli* forms a septum during division and *C. crescentus* does not, the OMs of both bacteria closely follow the PG layer during constriction. The diameter of OM tubes at the pre-separation stage, before complete daughter-cell division, is labeled for each species.

Cells divide their PG using a well-organized PG remodeling machinery called the divisome although the mechanisms driving constriction can differ^7^. For example, *E. coli* separates daughter cells by forming a septum PG perpendicular to the cell’s long axis^8–10^, whereas *Caulobacter crescentus* “pinches” PG inward at the division site without producing a septum^11^ (**Fig. 1**). In contrast, the mechanisms underlying OM division remain less understood, as the OMs of Gram-negative bacteria are devoid of energy sources and therefore lack dedicated division machinery.

Many Gram-negative bacteria tether their OMs to PG. Their physical linkage makes the periplasm a mechanical unit that balances turgor across the inner membrane^6^. For instance, *E. coli* employs multiple systems simultaneously to tether the OM to PG (**Fig. 1**). First, the PG-associated lipoprotein (Pal) that attaches to PG noncovalently and connects to the OM components in the Tol-Pal system, accumulates at the division site^12,13^. Second, the OM contains abundant Braun’s lipoprotein (Lpp) that attaches to PG directly through covalent bonds^14^. Unlike Pal, Lpp is absent from the division sites in wild-type *E. coli*. However, cells lacking both Pal and Lpp cannot separate their OMs during division, suggesting that Lpp can functionally compensate for the loss of Pal at division sites^15^. Third, some β-barrel OM proteins (OMPs) interact with PG noncovalently through their C-terminal OmpA-like domains^16^. Through these tethering mechanisms, the *E. coli* OM closely follows PG during constriction, forming a deep V-shaped invagination^8,17^ (**Fig. 1**). As the divisome assembles the septum toward the center of cell, the distance between the OM and PG remains remarkably consistent (**Fig. 1**)^9,10^, reflecting their tight coordination during cell division. *C. crescentus* does not produce Lpp but does encode Pal and OmpA-like proteins. During the gradual constriction of *C. crescentus*, its OMs remain attached to the PG layer until just before PG fission. At that stage, the PG separates first, leaving the two daughter cells connected by a narrow OM bridge, sometimes as small as 60 nm in diameter^11^ (**Fig. 1**).

In both *E. coli* and *C. crescentus*, how OMs separate after PG fission is less understood. It is unclear if the coordinated constriction of PG and OM is a universal division mechanism among Gram-negative bacteria and if the connection between OM and the divisome is essential for OM division. To answer these questions, we investigated cell division in *Myxococcus xanthus*, a rod-shaped Gram-negative bacterium^18^. In contrast to the rigid *E. coli* OM, the *M. xanthus* OM appears highly deformable^19^. Using bright field microscopy, *in situ* cryogenic electron microscopy (cryoEM), cryogenic electron tomography (cryoET) and fluorescence microscopy, we found a large disconnection between the PG and OM during *M. xanthus* division. Here, the OMs do not closely follow the constriction of PG during division. Instead, the OMs only invaginate slightly and remain largely unconstricted even after PG fission is complete, thus separating substantially later than the PG layer. However, reinforcing the OM-PG connection by overexpressing the endogenous tethering protein Pal synchronizes the constriction of both layers. We propose that the OMs separate by mechanical forces after PG fission and that the diversity of OM dynamics during division reflects a spectrum, where each organism’s OM position is determined by the strength of its PG-OM interactions.

## Results

### *M. xanthus* daughter cells remain transiently connected through their OMs after PG fission

*M. xanthus* lacks flagella and cannot swim. Instead, it moves across surfaces using two distinct mechanisms: a twitching-like motility driven by type IV pili (T4P) and a gliding motility powered by fluid motor complexes^20,21^. Cells were grown in a nutrient-rich liquid medium, subsequently spotted onto agar pads, and imaged using differential interference contrast (DIC) microscopy. In comparison to the wild-type, the nonmotile mutant lacking both T4P and a functional gliding motor (*ΔaglR pilA⁻*)^22,23^, frequently produced elongated cells and displayed increased heterogeneity in cell length (**Fig. 2a, S1**). To further investigate how motility systems affect cell length on solid surfaces, we quantified cell length of the wild-type that move by both motility systems and the mutants that move by either gliding (*pilA⁻*) or T4P-dependent motility (*ΔaglR*). In contrast to the nonmotile (*ΔaglR pilA⁻*) cells, all three motile strains displayed comparable and homogenous cell length (**Fig. 2a, S1**). We noticed that the loss of motility did not affect the length of individual (single) cells of the nonmotile strain but instead led to the formation of many cells approximately twice the length of a single cell (double, **Fig. 2a, S1**). Therefore, the length heterogeneity among nonmotile cells arose from delayed separation of dividing cells, rather than impaired septal PG synthesis. We reason that most dividing cells separated normally in liquid media and the cells of double length are the ones that were yet to separate on agar surfaces. Consistent with this hypothesis, when cells were fixed with formaldehyde in liquid medium prior to imaging, all three motile strains exhibited pronounced cell-length heterogeneity comparable to the nonmotile cells (**Fig. 2a**). These observations suggest that whereas the shear forces in shaking liquid culture can separate daughter cells, completion of daughter separation on solid surfaces depends on motility. Importantly, either motility system is sufficient for daughter cell separation on surfaces. For simplicity, we decided to use the wild-type cells that move by both motility systems to further study daughter cell separation in real-time using bright field microscopy (**Fig. 2a, b**).

**Fig. 2.**
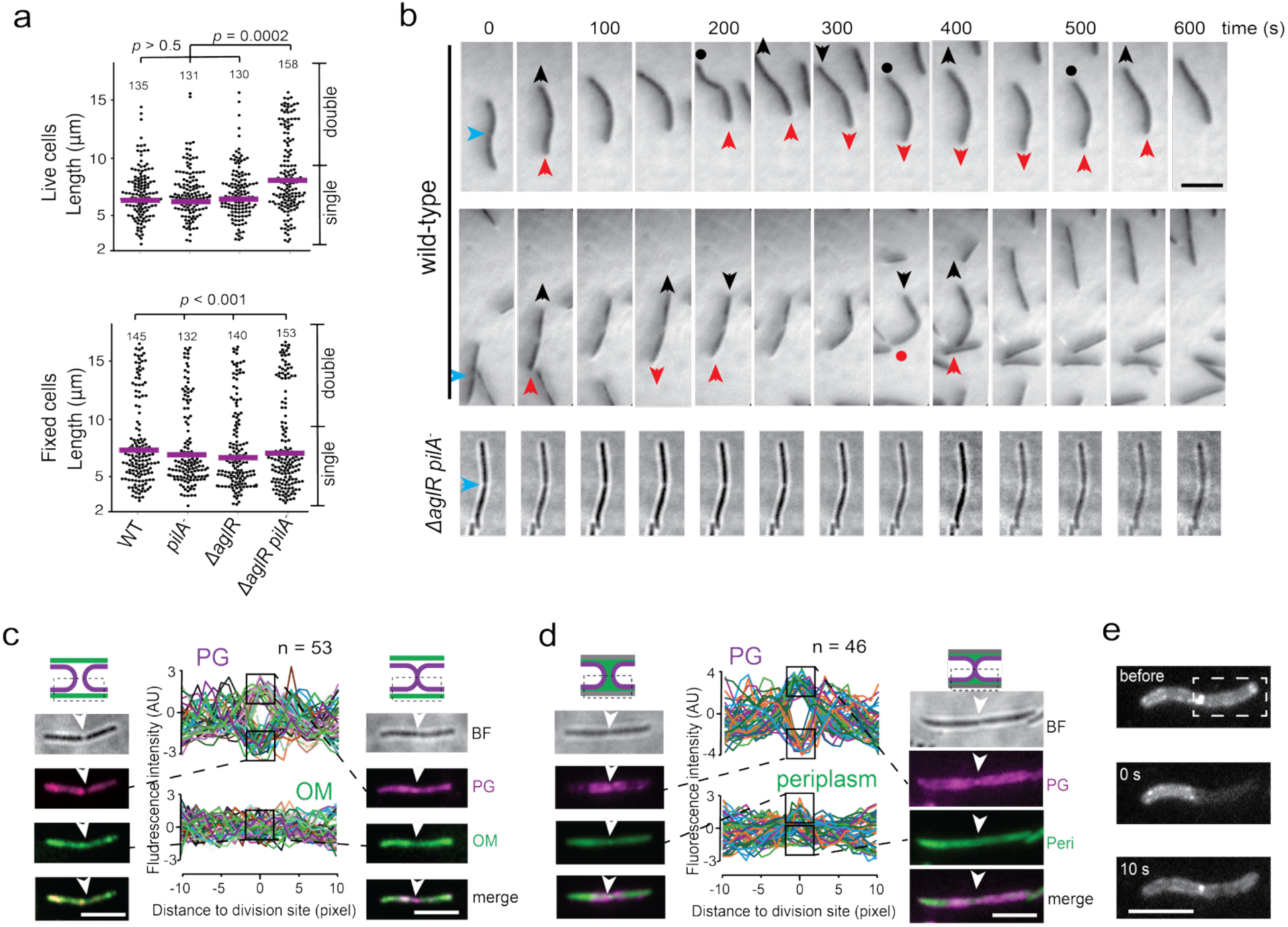
*M. xanthus* OMs separate after PG fission. **a)** On agar pads, live nonmotile cells (*ΔaglR pilA⁻*) were frequently elongated and displayed increased heterogeneity in length, whereas this phenotype was not observed with motile cells (wild-type, Δ*aglR*, and *pilA⁻*). In contrast, all four strains exhibited comparable cell-length heterogeneity when fixed in liquid culture and imaged on agar surfaces. The length ranges of single cells and unseparated sister cells (double) are marked on the right. For each strain, the number of measured cells is shown above the plot. Purple bars indicate the mean. *p* values were calculated using a single-tailed ANOVA test on unweighted samples. **b)** Nascent daughter cells of the motile wild-type cells move in an asynchronized manner before daughter cell separation (also see **Movies S1 – S6**). Black and red arrows indicate the moving directions of daughter cells, the blue ones point to the pale zones at the division sites, and the black and red dots indicate the frames in which cells’ movements stall. In contrast, the nonmotile Δ*aglR pilA⁻* cells were unable to complete separation on agar surfaces during observation periods (also see **Movies S9, S10**). **c, d)** Whereas the fluorescence signals of TADA-labeled PG either peak or plunge at the division sites, corresponding to septum assembly and PG fission, respectively, the WGA-labeled OMs (**c**) and msfGFP-labeled AlkP, a periplasmic protein (**d**), remain connected. Images of typical pairs of daughter cells are shown where white arrows indicate division sites. The fluorescence intensities of PG, OMs, and periplasms from multiple cells are shown on the right. In both panels, the sections of cells imaged using our setup are marked by boxes of dashed lines and the identities of cells are distinguished by colors. The fluorescence intensity values of multiple cells (sample size n is shown above each plot) were normalized using Z-score standardization. **e)** FRAP of AlkP-msfGFP showing that the nascent daughter cells share a periplasm. The white box indicates the photobleached region. Scale bars, 5 μm.

Using DIC microscopy to image cell division on agar surfaces, we frequently detected regions of reduced cell mass at the centers of elongated cells, which appeared as pale zones (**Fig. 2b, S1**), reminiscent of the enlarged periplasmic spaces in plasmolyzed *E. coli*, where the IMs detaches from the OMs^5,24,25^. This observation suggests that, after PG separation, daughter cells remained connected, most likely via the OM. The nascent daughter cells exhibited asynchronous movements driven by reciprocal pushing and pulling forces. They moved either in the same direction or in opposite directions, transiently increasing or decreasing their separation distance, while intermittent pauses of one daughter cell were frequently observed. Despite these movements, the daughter cells remained connected (**Fig. 2b, Movies S1 – S6**). The *pilA^-^* cells that moved only with gliding motility exhibited similar behavior (**Movies S7, S8**). The maximum distance between the poles of wild-type daughter cells prior to OM separation was 0.79 ± 0.19 µm (*n* = 63). Treating the OMs of *M. xanthus* as thin elastic shells, we estimated their stiffness (Young’s modulus) to be at the level of 0.2 MPa. Reported values for *E. coli* cell-envelope stiffness range from 0.8 MPa^5^ to 2 MPa^26^, with one study estimating that the OM contributes stiffness comparable to that of the PG layer^5^. Relative to these measurements, the *M. xanthus* OM is estimated to be softer than the *E. coli* OM^5^. Young’s modulus (*E*) was calculated as

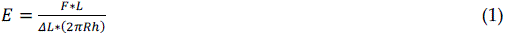

where *F* is the force generated by cell motility (∼200 pN. The T4P-dependent motility produces approximately 150 pN, whereas gliding motility contributes 5 – 50 pN^27^); *L* is the initial length of a dividing cell (∼10 µm), *ΔL* is the maximum OM extension (∼1 µm), *R* is the cell radius (∼ 0.5 µm), and *h* is the OM thickness (∼ 5 nm).

Using the same setup, isolated *ΔaglR pilA⁻* cells that lack motility did not exhibit any cell separation events on agar surfaces under time-lapse microscopy (**Fig. 2b, Movies S9, S10)**. Taken together, on solid surfaces, the natural niches of *M. xanthus*^18,20^, motility provides the mechanical force for daughter cell separation during the division of isolated cells.

To test the hypothesis that nascent daughter cells connect to each other transiently through OMs, we proceeded to image PG and OM fission of wild-type *M. xanthus* cells using fluorescence microscopy. Wheat germ agglutinin (WGA), a plant protein of 36 kDa, was reported to specifically bind to the OMs of *E. coli*^28^. To access WGA binding specificity in *M. xanthus* cells, we stained the entire OMs of cells expressing mCherry-labeled AglR, an IM protein part of a proton channel^29^, with Alexa Fluor 488-conjugated WGA for 1 h, then transferred cells to water. After this hypoosmotic shock, cells were plasmolyzed, i.e. the IMs separated from OMs^6,30^. While the IMs retained their original shape, the OMs frequently formed large bulges, which were specifically stained by WGA (**Fig. S2a**). WGA binds N-acetylglucosamine^31^, a sugar component in both the OMs and PG. To test if WGA penetrates the OMs and thus stains the PG, we grew cells overnight in the presence of TAMRA 3-amino-D-alanine (TADA) before WGA staining. TADA is a red fluorescent D-amino acid that is incorporated into PG by PG synthases^32^. After hypoosmotic shock, WGA and TADA displayed distinct labeling patterns in plasmolyzed cells (**Fig. S2b**). Therefore, the large size of WGA likely prevents it from crossing the OM barrier and binding to PG. Taken together, WGA can be used to distinguish OMs from IMs and PG *M. xanthus*.

To visualize PG and OMs simultaneously during division, we grew cells with TADA overnight then stained their OMs with WGA for 1 h. We selected dividing cells with cytoplasms separated by fewer than 3 pixels (480 nm) in DIC images and used fluorescence microscopy to visualize a ∼200-nm-thick longitudinal section encompassing both the PG and OMs. Imaging was performed under highly inclined and laminated optical sheet (HILO) illumination^23,29,33,34^. Under these imaging conditions, fluorescence signals from both OM and PG covered the cylindrical cell surfaces facing the coverslip instead of forming well-defined outlines around the cells. Fluorescence signals of PG at division sites reflected the progression of PG fission: intensified signals indicated septal PG growth (**Fig. 2c**, right), whereas loss of fluorescence signified complete PG separation (**Fig. 2c**, left). Regardless of the status of PG, all the dividing cells (n > 100) displayed continuous OM at their division sites (**Fig. 2c**). Crucially, we never detected a surge in OM signal at division sites, suggesting that, unlike PG, *M. xanthus* OMs do not undergo steep invagination (**Fig. 2c**). These distinct staining patterns further confirmed that WGA did not stain PG. Therefore, nascent daughter cells are still transiently connected by OMs after PG fission.

The continuity of the OMs during division suggests that nascent daughter cells share continuous periplasms before OM fission. To investigate this possibility, we labeled the periplasmic alkaline phosphatase AlkP (K1515_31585/MXAN_1389)^35^ with monomeric super-folder green fluorescent protein (msfGFP) in the wild-type background. AlkP-msfGFP, expressed ectopically from a phage attachment site under the control of the *pilA* promoter, displayed a uniform distribution throughout the cell envelope. Notably, AlkP-msfGFP was also detected within OM tubes that are not visible in bright-field microscopy^36^, further confirming its periplasmic localization (**Fig. S3**). During cell division, AlkP-msfGFP fluorescence remained continuous across the division site, regardless of whether PG had separated. Unlike the OM fluorescence, which was confined to the cell surface, AlkP signals showed an increase between daughter cells following PG separation, suggesting the presence of a continuous and relatively deep periplasmic compartment (**Fig. 2d**). In nascent daughter-cell pairs exhibiting a bright fluorescence focus between the cells, one daughter cell together with the intervening focus was photobleached (**Fig. 2e**). Following photobleaching, fluorescence from the unbleached daughter cell diffused into the bleached cell and re-established the bright focus (**Fig. 2e**). The rapid fluorescence recovery after photobleaching (**FRAP**) is not likely due to new protein synthesis. This result provides further evidence that nascent daughter cells share a continuous periplasm prior to separation. Together, these observations indicate that OM separation occurs substantially later than PG fission during *M. xanthus* cell division.

### OM and PG do not invaginate in a synchronized manner during *M. xanthus* cell division

To investigate OM fission at high resolution, we captured micrographs of wild-type *M. xanthus* cells using cryoEM, in which the electron densities of the IMs and OMs were easily distinguished from the background (**Fig. 3**). In contrast, the thin PG layer was extremely difficult to visualize (**Fig. 3**). In this case, we use IMs to proximate PG fission. Due to the long generation time of *M. xanthus* (∼4 h)^37^, locating individual dividing cells on EM grids was difficult. After screening more than 1,000 cells, we identified 36 at the constriction stage (**Fig. 3a–3c, S3, Table S1**) and 25 pairs of daughter cells that had completed PG fission (**Fig. 3d–3f, S3, Table S1**). Among the 36 constricting cells, the IMs exhibited varying degrees of invagination, whereas the OMs only constricted slightly, regardless of the extent of IM invagination (**Fig. 3a–3c, S3, Table S1**). Consequently, we did not observe any deep V-shaped OM invaginations, a feature commonly seen in dividing *E. coli* cells^8^. Among the 25 daughter cell pairs that had completed PG fission, 23 remained connected by the OMs of the mother cells (**Fig. 3d–3f, S3, Table S1**). We further confirmed the asynchrony between IMs and OMs using cryogenic electron tomography (**cryoET**) on dividing cells that displayed separated IMs but connected OMs in cryoEM images. As expected, the tomograms showed that OMs of these cells do not follow the IMs during constriction (**Fig. S4a**); they remained connected after IM separation (**Fig. 3g, S4b, Table S1**). Taken together, these observations suggest that OM and PG do not invaginate in a synchronized manner during *M. xanthus* cell division.

**Fig. 3.**
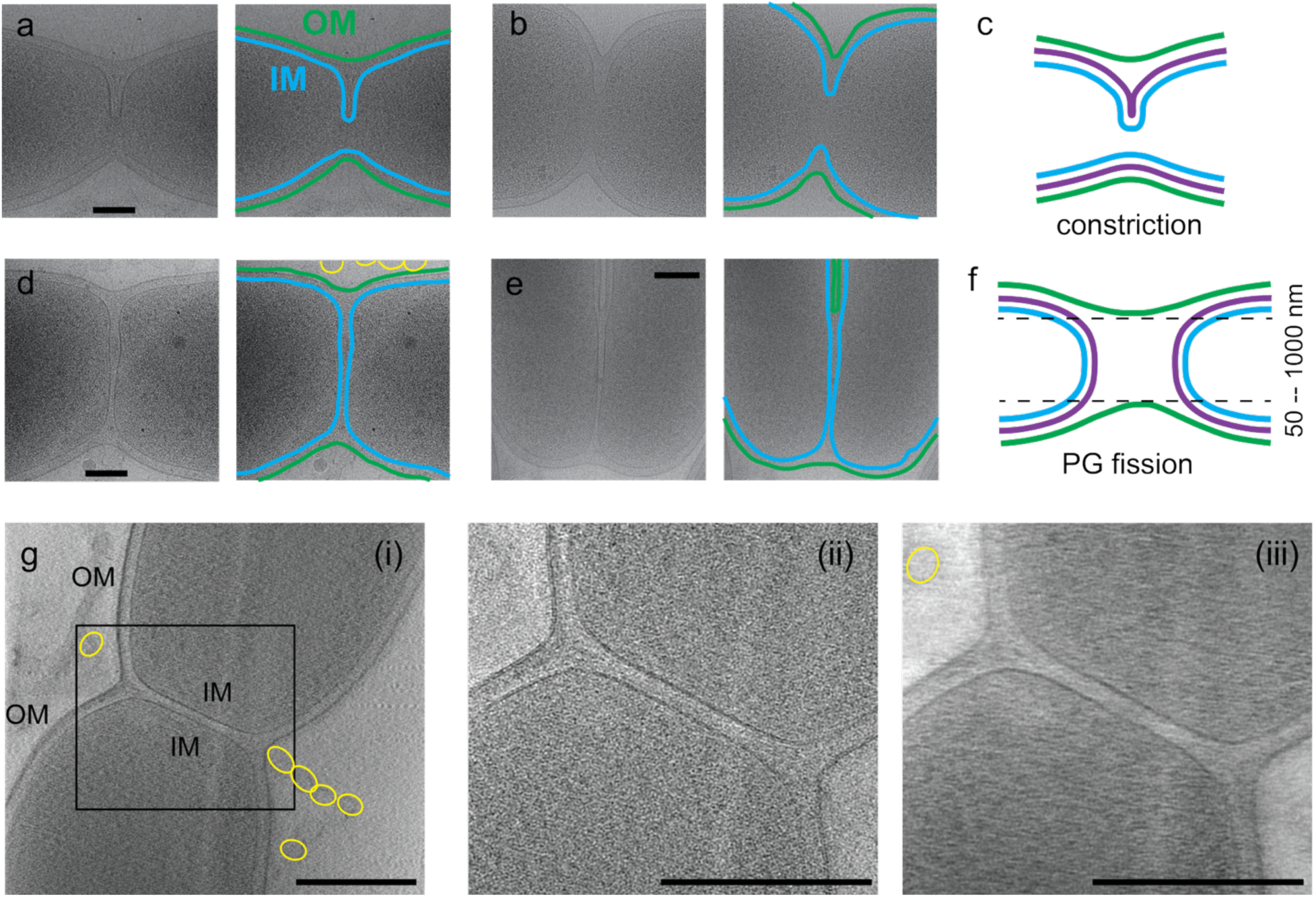
OM and PG do not invaginate in a synchronized manner during *M. xanthus* cell division. **a**, **b**) CryoEM images of cells at the stage of constriction. **c)** Schematic depiction of the PG constriction stage. **d, e**) CryoEM images of cells at the stage of PG fission. **f**) Schematic depiction of the PG fission stage. **g**) CryoET images of a cell at the stage of PG fission. (**i**) A representative slice from a 3D tomographic reconstruction of a dividing *M. xanthus* cell. (**ii**) Magnified view of the region highlighted in (**i**). (**iii**) Corresponding 40-nm-thick projection of the same region. Yellow circles mark extracellular vesicles frequently observed around *M. xanthus* cells. Scale bars, 200 nm.

Consistent with the asynchronized movements observed through DIC microscopy (**Fig. 2b**), the distance between the IMs at the new cell poles ranged from approximately 20 nm (**Fig. 3d–3f, S3, Table S1**) to about 500 nm (**Fig. 4a, 4b, S5, Table S1**). Consequently, their OMs were either relaxed (**Fig. 3d, S3**), stretched (**Fig. 4a, S5**) or folded on one side and stretched on the other (**Fig. 3e, S3**), and even twisted (**Fig. 4b**). Strikingly different from dividing *C. crescentus* cells^11^, narrow IM tubes were never visualized between daughter cells. Of the 25 daughter cell pairs that had completed PG fission, three exhibited markedly stretched OMs (**Fig. 4a, 4b, S5, Table S1**). The stretch of OMs indicates that *M. xanthus* OMs are highly elastic. This elasticity suggests that OMs might separate through a mechanical stretch-and-snap-back mechanism after PG fission. CryoEM imaging allowed us to capture two instances of stretched and ruptured OMs between separating daughter cells (**Fig. 4c, S5, Table S1**). In contrast, such OM ruptures were never observed on nondividing cells. Interestingly, OM bulges, the common indicators of the detachment between OM and PG^6^, were observed at sites where OMs were stretched (**Fig. 4b, c, Table S1**).

**Fig. 4.**
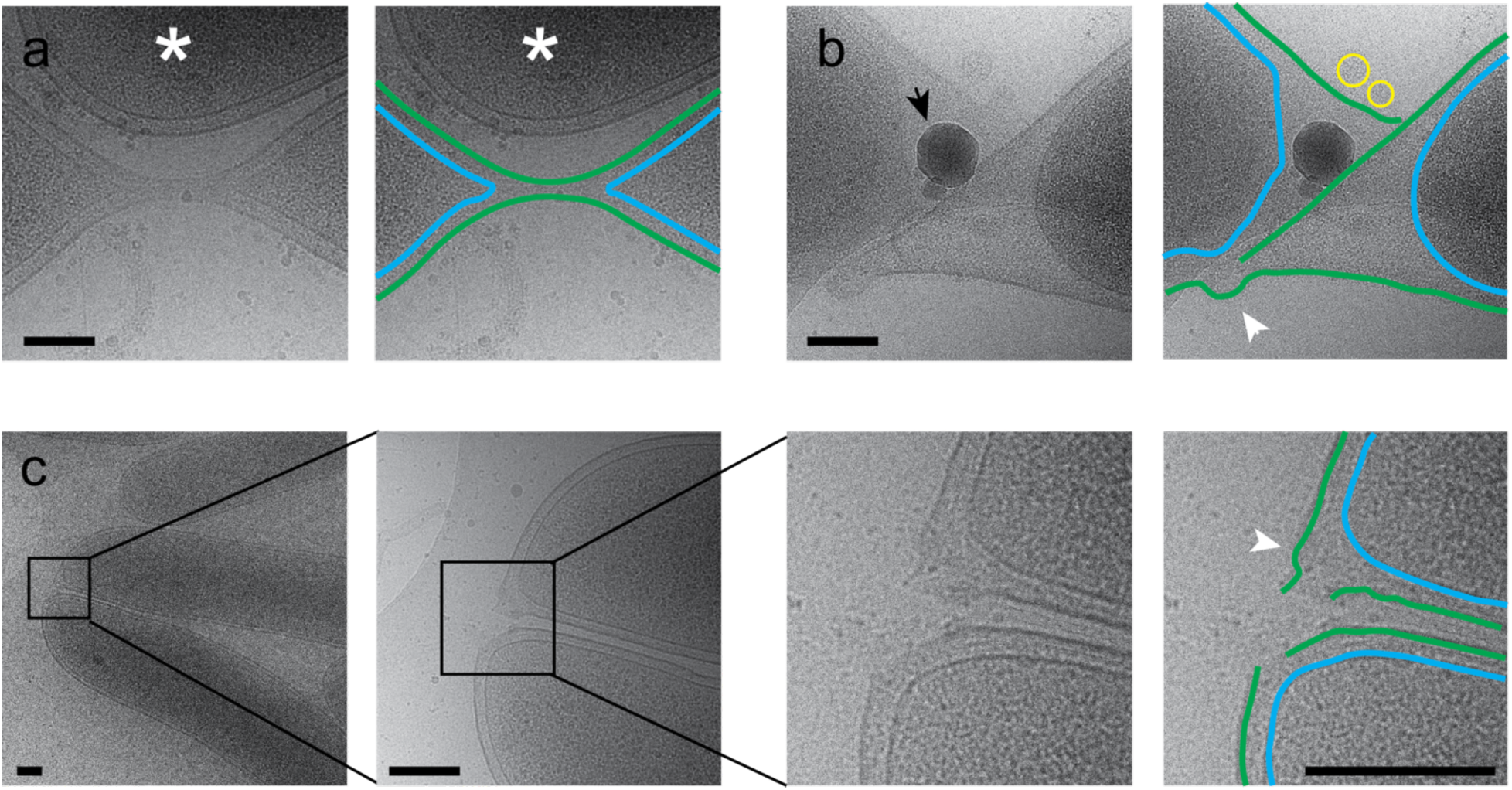
Stretching and subsequent rupture of the OMs indicate a physical mechanism underlying OM separation following PG fission. **a)** A pair of nascent daughter cells connected by a stretched OM. The Asterisk marks a third cell located near the separating daughter cells. **(a)** and stretched and twisted (**b**) OMs. **c)** The rupture of OMs. White arrows point to bulges on the OMs while the black one points an ice particle. Yellow circles mark extracellular vesicles. Scale bars, 200 nm.

**Fig. 5.**
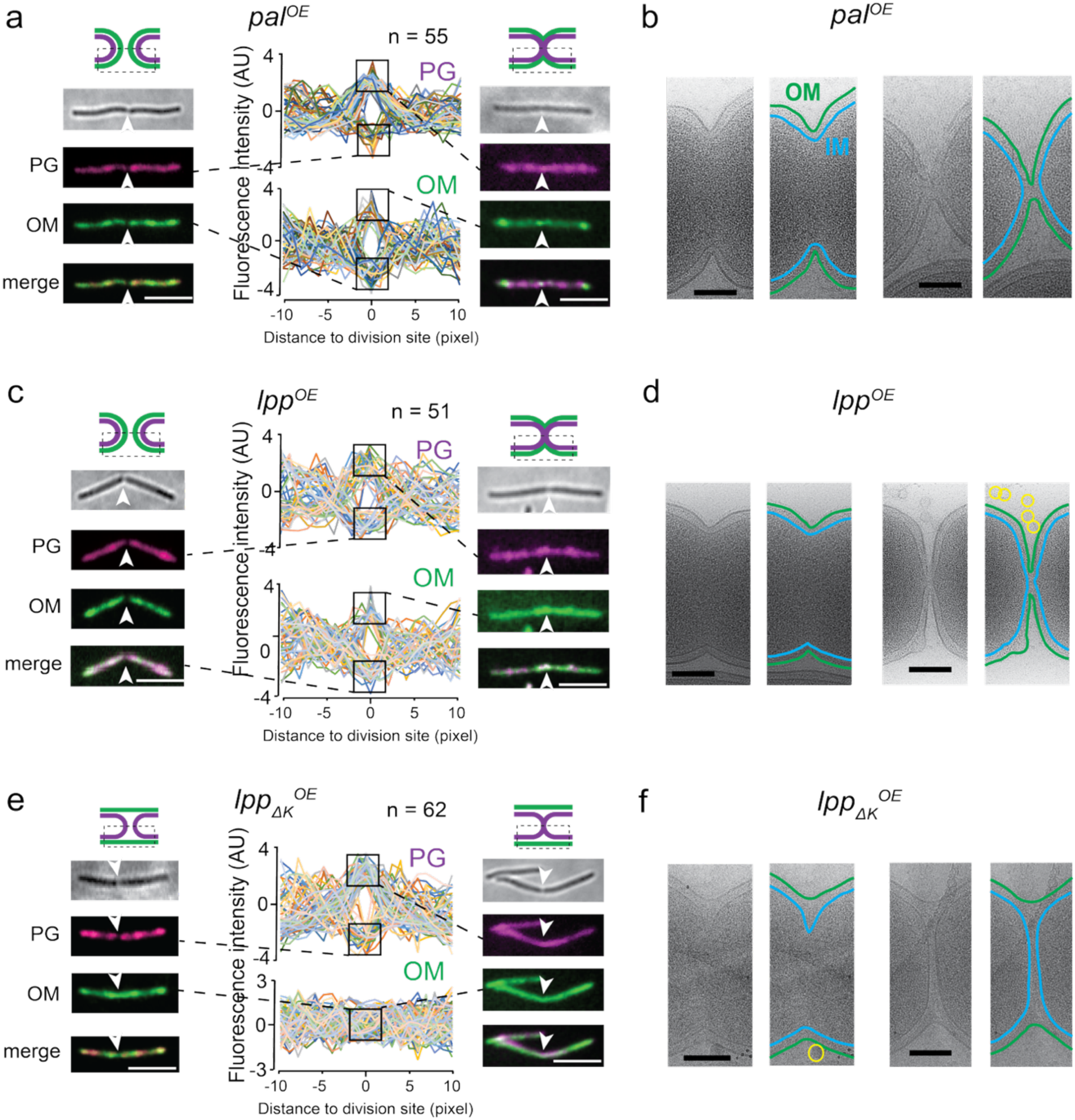
Enhanced OM-PG connection synchronizes the constriction of both layers. **a**, **c**, **e**) Fluorescence images of typical dividing cells that overexpress native Pal (**a**), wild-type *E. coli* Lpp (**c**) and an *E. coli* Lpp variant that lacks the lysine residue that attaches OM to PG (Lpp_ΔK_, **e**). The PG and OMs of cells are labeled with TADA and Alexa Fluor 488-conjugated WGA, respectively. White arrows point to the division sites. The fluorescence intensities of PG and OMs from 12 cells are shown on the right. The sections of cells imaged using our setup are marked by boxes of dashed lines and the identities of cells are distinguished by colors, and the fluorescence intensity values were normalized using Z-score standardization. **b, d, f)** representative cryoEM images of the division sites in cells at the constriction (left) and PG fission (right) stages. Yellow circles mark extracellular vesicles. Scale bars, a, c, e, 5 μm, b, d, f, 200 nm.

Strikingly, the stretched OMs frequently deformed the nascent cell poles at division sites, suggesting that the PG at the new poles is remarkably soft (**Fig. 4, S5**). This combined flexibility in both the OM and PG layers may be a general property of *M. xanthus* PG throughout the cell body, as cells often bend into “S” and “U” shapes on surfaces (**Fig. S7**).

Another unexpected finding is that the IMs of all the 36 constricting cells invaginated in an asymmetric pattern, suggesting asymmetric septum growth (**Fig. 3a–3c, S4a**). The absence of synchronized constriction between OMs and IMs is strikingly similar to an *E. coli Δlpp Δpal* strain that loses the major connections between PG and OM^15^, producing an OM invagination defect as reported here with WT *M. xanthus* cells. However, different from the *E. coli* mutant, wild-type *M. xanthus* cells do not form cell chains, indicating that the uncoupled invagination of OMs and IMs does not lead to division defects.

To determine whether asymmetric cell constriction is a common feature of dividing *M. xanthus* cells, we imaged wild-type cells following a 10-min pulse labeling with TADA. In cells approximately twice the length of typical vegetative cells, TADA foci were frequently observed at the cell midpoint, indicating PG growth at future division sites. Notably, TADA signals at these sites often exhibited pronounced asymmetry across long cell axes. In some cells, a single TADA focus was detected on one side of the division site, whereas in others, two foci were present on opposite sides, with one focus consistently brighter than the other (**Fig. S8a, b**). Because septal PG growth is coordinated by a constricting ring formed by the tubulin homolog FtsZ^38^, such asymmetric septal PG growth suggests that the FtsZ cytokinetic rings may position asymmetrically. We investigated the localization of the FtsZ ring in the wild-type background. As fluorescence labeled FtsZ variants do not support *M. xanthus* growth in the absence of native FtsZ^38^, we used the plasmid pKA51^38^ to construct a merodiploid *ftsZ^+^/ftsZ-gfp* strain that expressed green fluorescent protein (GFP)-labeled FtsZ using its native promoter at the Mx8 phage attachment site. Consistent with a previous report^38^, FtsZ-GFP displayed dynamic localization that changed over time from diffuse/speckled to bright clusters near the longitudinal center of many cells, marking the assembly of the cytokinetic Z-ring. FtsZ clusters in these cells always positioned asymmetrically, offset from their long cell axes (**Fig. S8c, d**). Therefore, consistent with TADA and cryoEM observations, the *M. xanthus* FtsZ rings constrict in an asymmetric manner.

### Enhanced OM-PG connections synchronize the constriction of both layers

Similar to most Gram-negative bacteria, *M. xanthus* possesses a conserved Tol-Pal system and several OmpA-like OMPs^35^, albeit cryoEM and fluorescence imaging demonstrated that their native expression fails to couple the OM to PG during division (**Fig. 2–4**). To examine how PG-tethering influences the mode of OM division, we overexpressed the native Pal (K1515_14780/MXAN_4581, 38.6% sequence identical to *E. coli* Pal)^35^ as a merodiploid in *M. xanthus* by inducing a vanillate-inducible promoter^39^ with 100 µM sodium vanillate. In the presence of excess Pal, OM fluorescence rose and fell in synchrony with PG fluorescence (**Fig. 5a**), suggesting tight coupling between these two layers during constriction. Consistently, cryoEM analysis revealed that OM constrictions tracked IMs closely in all 29 dividing cells observed (**Fig. 5b, S3, Table S1**), a pattern not observed in the wild-type *M. xanthus*, but in dividing *E. coli*^8^. Although *M. xanthus* lacks Lpp-like proteins, it encodes two putative L,D-transpeptidases (K1515_32695/MXAN_1173 and K1515_09345/MXAN_5692)^35^, the enzymes that form L,D-crosslinks in the PG in many bacteria^40^ and catalyze the covalent linkage between the terminal lysine residue of Lpp and the stem peptides of PG in *E. coli*^41^. To test if introducing *E. coli* Lpp into *M. xanthus* could strengthen the OM-PG connection and thus synchronize their constriction during cell division, we used the same vanillate-inducible promoter to express wild-type *E. coli* Lpp exogenously. Remarkably, like Pal, overexpressed Lpp synchronized OM and PG constriction in fluorescence images (**Fig. 5c**) and OM and IM constriction in all 26 cryoEM micrographs (**Fig. 5d, S3, Table S1**). These observations suggest that exogenous Lpp strengthens the OM-PG connection, enabling OMs to constrict in concert with PG and IMs at division sites. To confirm that the coupling effect of Lpp is based on its covalent linkage to the PG layer, we overexpressed an Lpp variant that lacks the terminal lysine residue (Lpp_ΔK_) and therefore is unable to form covalent bonds with PG^41^. Overexpressed Lpp_ΔK_ failed to synchronize OMs with either PG (**Fig. 5e**) or IMs (**Fig. 5f, S3, Table S1**). Thus, reinforcing the OM-PG connection, either by endogenous or exogenous coupling proteins, is sufficient to synchronize their constriction during cell division.

Cells overexpressing Pal and Lpp displayed symmetric IM invaginations (**Fig. 5b, 5d, S3, Table S1**), in stark contrast to the asymmetric invaginations observed in the wild type cells (**Fig. 3a, 3b, S3**) and the ones overexpressing Lpp_ΔK_ (**Fig. 5e, 5f, S3, Table S1**). To quantify the asymmetry of IM invagination, we calculated the ratio of OM–IM distances at the leading and lagging edges. For the wild type and Lpp_ΔK_ –expressing cells, the asymmetric ratios are 2.94 ± 1.39 (n = 36) and 3.23 ± 0.93 (n = 18), respectively. In contrast, the asymmetric ratios for the cells expressing Pal and wild-type Lpp are 1.16 ± 0.08 (n = 26) and 1.17 ± 0.06 (n = 19), respectively (**Fig. S9**). These results suggest that although *M. xanthus* lacks the Lpo-like proteins that directly activate PG-assembly enzymes^17,42–44^, other OM-PG coupling proteins can also regulate septum PG growth.

Whereas Lpp partially rescues division defects caused by the absence of Pal, Lpp does not localize to division sites in wild-type *E. coli* cells^15^. To test whether exogenous Lpp accumulates to the division sites of *M. xanthus*, we performed immunofluorescence microscopy using antibodies against *E. coli* Lpp. The antibodies bound relatively uniformly across the surface of *E. coli* MG1655 cells but were largely excluded from division sites (**Fig. S10**). In contrast, they showed no detectable binding to wild-type *M. xanthus* cells, thereby confirming their specificity. In *M. xanthus* cells expressing wild-type Lpp, Lpp distributed throughout the cell envelope and became enriched at division sites and newly formed poles in dividing cells. Whereas Lpp_ΔK_ showed similar homogeneous distribution, it did not exhibit accumulation at division sites (**Fig. S10**). Thus, Lpp accumulates at division sites through its covalent attachment to PG, although the mechanism underlying this accumulation remains unclear.

## Discussion

OM fission is the final, yet often overlooked, step of Gram-negative bacterial cell division. It is tempting to presume that if the OMs simply follow PG invagination, PG separation will naturally sever the OMs between daughter cells. However, the synchrony between OM and PG often diminishes before OM fission, indicating that OM separation cannot be explained solely as a consequence of OM-PG coupling. For instance, during *E. coli* division, the OM invaginates in concert with the PG layer throughout most of the constriction process^8^. However, following septal PG separation, the OM remains connected by an aperture approximately 200 nm in diameter that must still be sealed (**Fig. 1**). What mechanism drives OM fission after the septal PG has split? Similarly, *C. crescentus* OMs follow PG invagination even deeper. After PG separation, cells remain connected only by OM tubes about 60 nm wide (**Fig. 1**)^11^. What force ultimately severs these OM tubes remains to be answered (**Fig. 1**)? *M. xanthus* provides a unique opportunity to examine this terminal step of Gram-negative cell division. In *M. xanthus*, OM division does not require the OMs to constrict in synchrony with the PG, demonstrating that tight PG-OM couplings are not essential for OM scission.

When a bacterial OM is approximated as a cylindrical tube, its stiffness depends on its elastic modulus and cross-sectional area^4^. In *E. coli* and *C. crescentus* with stiff OMs, tight OM-PG coupling and thus synchronous invagination, likely play a critical role in OM fission by narrowing the OM necks to about 200 nm and 60 nm wide, respectively, and thereby reducing the force required for membrane separation (**Fig. 1**). Whereas OM necks still undergo narrowing during *M. xanthus* daughter cell separation (**Fig. 4**), their soft OMs allow cell separation to proceed with relatively little force, making PG-tethered OM invagination less critical. We proposed that in the last step of division, mechanical forces generated by shear stress, Brownian motion in liquid, cell motility, or cell–cell collisions, may drive OM separation between daughter cells. However, can OM fission in organisms with relatively stiff OMs also occur through a purely physical mechanism?

How OMs rupture, reseal, and reconnect with the PG layer remains unclear. Rayleigh–Plateau instability, which describes the surface-tension–driven breakup of cylindrical fluid columns to minimize interfacial energy, offers a useful conceptual framework for interpreting this process, as illustrated in liquid jet fragmentation in air and bubble pinch-off in liquid^45,46^. The OMs of *M. xanthus* meet several conditions that could permit a Rayleigh–Plateau–like instability: (i) a cylindrical geometry; (ii) high deformability; (iii) a scale (∼1 µm radius) at which membrane tension may exceed bending rigidity; and (iv) weak OM–PG coupling, which minimizes external mechanical constraints. Consistent with this framework, the observed thinning of OM tethers between nascent daughter cells resembles the initial stages of capillary instability in a membrane cylinder. These observations suggest that analogous to liquid jet fragmentation and bubble pinch-off, OMs may spontaneously reseal following fission. Supporting this idea, the OM bulges observed both immediately before (**Fig. 4b**) and after (**Fig. 4c**) fission may represent manifestations of such a resealing process^45,46^. Because separation driven by Rayleigh–Plateau–like instability typically occur on microsecond timescales^45,46^, OM fission is likely an extremely rapid process. Given the resolution of light microscopy and the length of the cell cycle of *M. xanthus* (∼4 h), it will be close to impossible to sustain acquisition at this frame rate over a period long enough to capture a division event and resolve stretch-and-snap-back behavior of the OMs. Nevertheless, tether-pulling experiments using optical tweezers or micropipette aspiration could be valuable for assessing cell integrity following OM rupture and for quantifying the longitudinal stiffness of the *M. xanthus* outer membrane.

Although the soft OMs of *M. xanthus* permit OM separation in the absence of strong OM-PG tethering, our results do not indicate that OM softness is mechanistically linked to OM-PG tethering. Rather, our findings demonstrate that bacteria can function with substantially different degrees of OM-PG tethering. For instance, although Lpp or Pal are important for OM stability in *E. coli* and their absence often causes OM to blebbing, neither protein is essential for cell division^47,48^. Conversely, reinforcing the OM-PG connection does not impede *M. xanthus* division. These observations suggest that the connections between PG and OMs are less conserved than the structures of these cell wall layers.

## Materials and methods

### Strains and growth conditions

Vegetative *M. xanthus* cells were grown in liquid CYE medium (10 mM MOPS pH 7.6, 1% (w/v) Bacto™ casitone (BD Biosciences), 0.5% yeast extract and 8 mM MgSO_4_) at 32 °C, in 125-ml flasks with vigorous shaking, or on CYE plates that contains 1.5% agar. We used strain DZ2 as the wild-type *M. xanthus* strain^49^. Overexpression mutants were constructed by electroporating DZ2 cells with 4 µg of plasmid DNA. The genes, driven by a vanillate-inducible promoter, were inserted into the Mx8 phage attachment site as merodiploid on the *M. xanthus* chromosome. Transformed cells were plated on CYE plates supplemented with 10 µg/ml tetracycline and expression was induced by 100 µM sodium vanillate. The strains, plasmids, and primers used in this study are listed in **Table S2**. Three technical replicates and three biological replicates were imaged for the published and newly constructed strains, respectively.

### Immunofluorescence and live cell Light microscopy

Cells from vegetative CYE cultures (0.5 ml) at OD_600_ 1 were fixed for 20 min after transfer to tubes containing 100 µl paraformaldehyde (16%) and 0.2 µl glutaraldehyde (25%). Fixed cells were then immobilized on freshly prepared poly L-lysine-treated slides and treated for 4 min with 1 µg/ml lysozyme treatment. Washing was performed in PBS and probing in PBS 2% BSA with the anti-Lpp antibodies (abbexa, catalog number abx345344) added at 1:100 dilution. Lpp was visualized using an Alexa Fluor™ Plus 488 conjugated goat anti-rabbit IgG (H+L) highly cross-adsorbed secondary antibody (ThermoFisher Scientific, catalog number A32731TR, 1:200).

For bright field and fluorescence microscopy on live cells, cultures were grown in liquid CYE overnight to OD_600_ ∼1 and stained by 75 µM TADA for 1 h. Then 10 µg/mL Alexa Fluor-488 conjugate WGA was added to the culture, and cells were grown for 1 h. Cells were spun down at 6,000 x g for 3 min, and the pellet washed three times with CYE. 5 µl of cells were spotted on agar (1.5%) pads that contain 8 mM MgSO_4_. Images were captured using a Andor iXon Ultra 897 EMCCD camera (effective pixel size 160 nm) on an inverted Nikon Eclipse-Ti™ microscope with a 100 ✕ 1.49 NA TIRF objective. Fluorescence of TADA and Alexa Fluor 488-conjugated WGA was excited by the 561-nm and 488-nm lasers, respectively. Fluorescence intensities were quantified using the ImageJ suite (https://imagej.net). To convert raw data points into a universal scale with a mean of 0 and a standard deviation of 1, we used Microsoft Excel to normalize fluorescence intensities using Z-score standardization, Z = (χ – µ)/σ, where Z, χ, µ, and σ are the standard score, observed value, mean, and standard deviation, respectively. For FRAP imaging, cells were photobleached using a 488-nm laser beam focused by a Nikon Ti-TIRF-PAU illuminator.

### CryoEM and cryoET

Bacterial strains were grown overnight in CYE media with appropriate antibiotics, incubated with shaking at 32 °C and 250 rpm to an optical density of OD_600_ 0.6. Cells were collected by centrifugation (3 min, 6,000 × g, 25 °C) and resuspended in CYE media to a final OD_600_ 12. This cell suspension (3 µl) was applied to C Flat-1.2/1.3 200 mesh copper grids (Electron Microscopy Sciences) that were glow discharged for 30 seconds at 15 mA. Grids were plunge-frozen in liquid ethane with an FEI Vitrobot Mark IV (Thermo Fisher Scientific) at 4 °C, 100% humidity with a waiting time of 30 s, two-side blotting time of 2.5 – 3 s, and blotting force of 0. All subsequent grid handling and transfers were performed in liquid nitrogen. Grids were clipped onto cryo-FIB autogrids (Thermo Fisher Scientific).

Images were acquired using a Thermo Fisher Scientific Titan Krios G4 transmission electron microscope, equipped with a Gatan K3 direct electron detector and a Gatan BioContinuum energy filter (pixel size 0.27 nm, except in the left panel of Fig. 3g, which is 2.06 nm). Imaging was performed using a 15 eV slit width on the energy filter to enhance image contrast. Micrographs were recorded in counted mode at a nominal magnification of 33,000x, using a dose rate of 14.65 electrons/pixel/second over a 2.6-second exposure, resulting in a total accumulated dose of 50 electrons/Å². CryoEM images were corrected for beam-induced motion and subjected only to cropping and contrast adjustments, with no other image processing performed.

CrytoET was performed as reported^50^. Data were collected on a Krios transmission electron microscope equipped with a slit energy filter. Tilt series were acquired at a nominal magnification of 33,000× with a defocus of approximately 3 µm, resulting in a calibrated pixel size of 0.27 nm/pixel. A total of 33 tilt images were collected over an angular range from −48° to +48° with 3° increments. At each tilt angle, images were recorded in dose-fractionation mode, generating six subframes per projection. Beam-induced motion was corrected using MotionCor2^51^. The corrected images were assembled into tilt-series stacks using an in-house script. Tilt-series alignment was performed with AreTomo to generate aligned stacks^52^. Three-dimensional tomographic reconstructions were generated using Tomo3D with the simultaneous iterative reconstruction technique (SIRT)^53^. The reconstructed tomograms were subsequently denoised using TomoAND^54^. Finally, three-dimensional visualization and analysis of the tomograms were performed using IMOD^55^.

## Acknowledgments

We thank Drs. Michael Van Nieuwenhze and Yen-Pang Hsu for providing TADA, Drs. Anke Treuner-Lange and Lotte Sogaard-Andersen for the pKA51 plasmid, and Dr. Gaya Yadav (Laboratory for Biomolecular Structure and Dynamics (LBSD)-CryoEM at TAMU) for the assistance on cryoEM imaging. Part of this work was supported by the National Institutes of Health grants GM129000 to B. N. and GM140886 to D.W. We received financial support from Dr. David R. Zusman, who played no role in the design, execution, or presentation of this work.

## Author Contribution Declaration

C.B.D.S. and B.N. conceived the project, designed the study, analyzed the data, and wrote the manuscript. C.B.D.S., C.A.R.C., S.Z. G.P.S., D.W. and B.N. performed the experiments. C.B.D.S., D.W. and B.N. revised the manuscript.

## Data Availability

All data are presented in the main text and supplementary materials.

