## supplements for "A Tether-Independent Mechanism for Bacterial Outer Membrane Separation"

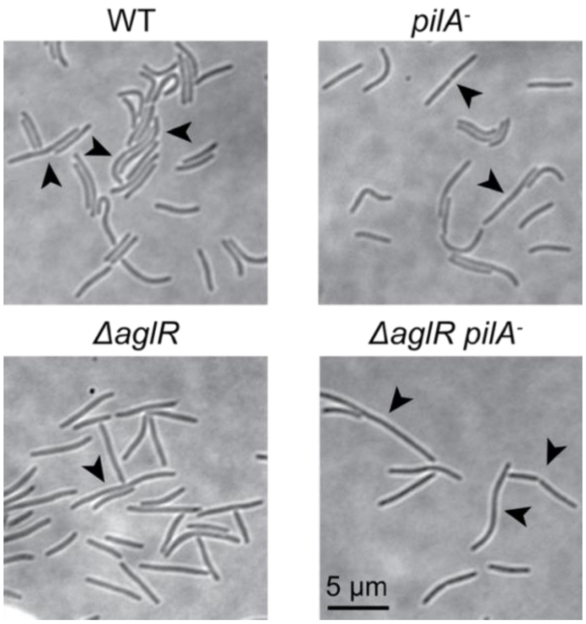


**Fig. S1. Cell morphology of the wild-type and motility mutants.** Black arrows point to the elongated cells that are potentially undergoing division. Cells were grown in liquid media, spotted, and visualized immediately.

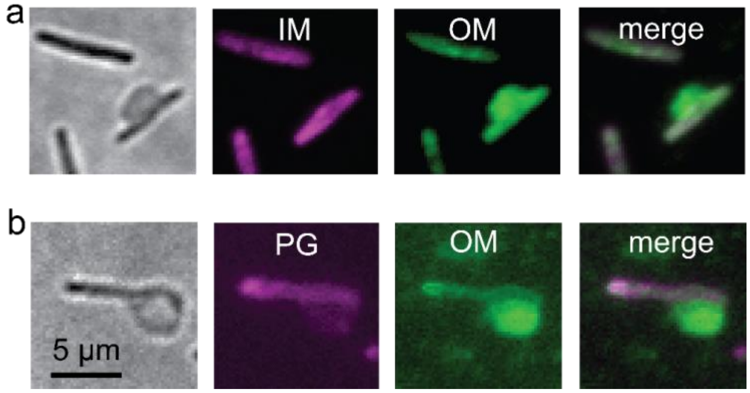


**Fig. S2. Plasmolysis assay indicates that WGA specifically label the OMs of *M. xanthus*.** OM bulges formed under hypoosmotic shock, which were specifically stained by Alexa Fluor 488-conjugated WGA. The labeling patterns of WGA are distinct from IM (**a**) and PG (**b**). The IMs and PG are labeled by AglR-mCherry and TADA, respectively.

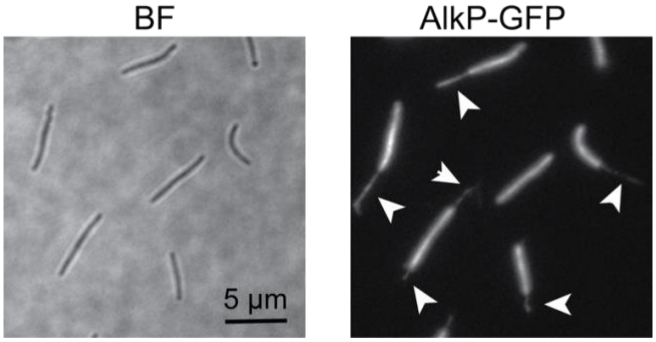


**Fig. S3. AlkP-GFP localizes to the periplasm and OM tubes of *M. xanthus*.** Arrows point to the OM tubes with periplasmic content.

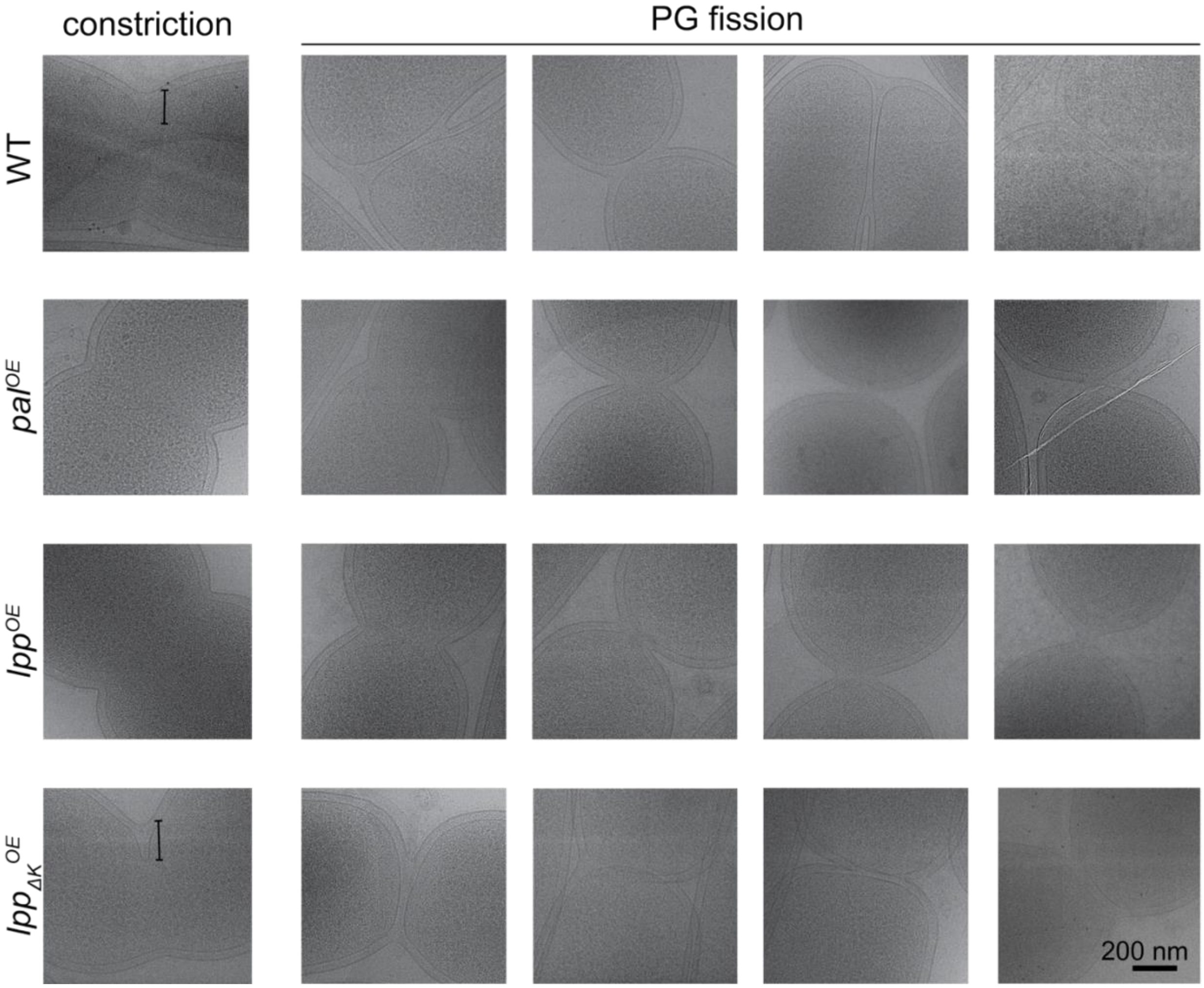


**Fig. S4. A gallery of cryoEM images of cells in the constriction and PG fission (marked by IM fission) stages.** Distance indicators mark the delayed invagination of OM in comparison to the PG.

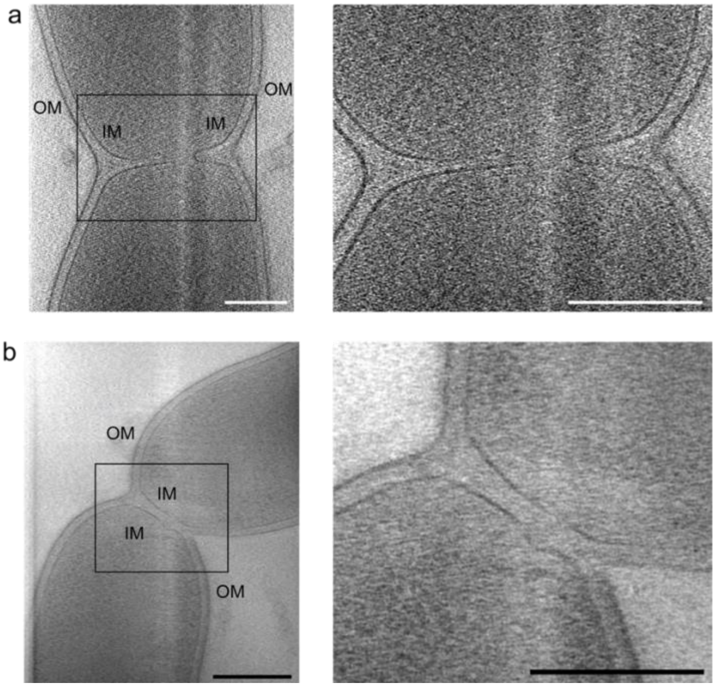


**Fig. S5. CryoET images of cells at the stages of constriction (a) and cytokinesis (b).** Left, representative slices from 3D tomographic reconstruction of dividing *M. xanthus* cells. Right, magnified views of the regions highlighted on the left. Scale bars, 200 nm.

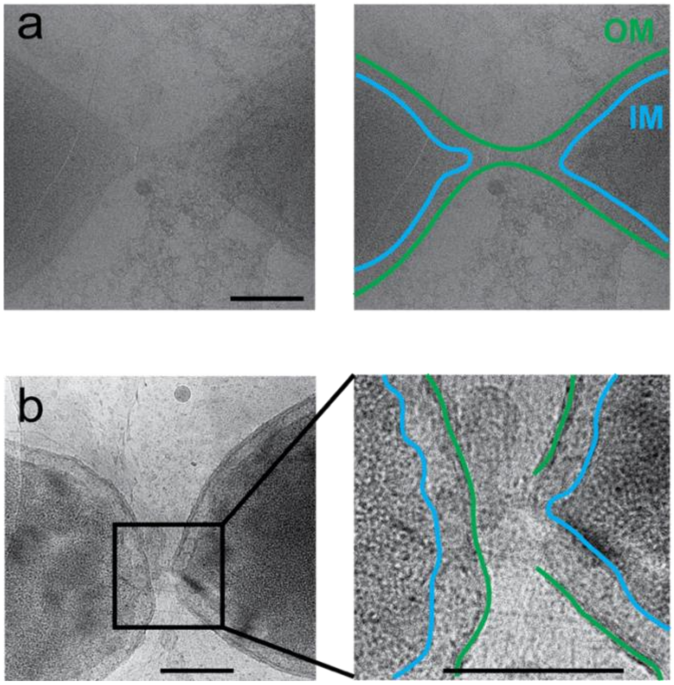


**Fig. S6. Additional examples of the stretch (a) and rupture (b) of OM.** Scale bars, 200 nm.

**Table S1. Summary of cryoEM observations on dividing cells.** For each strain, the number of cell pairs on each division stage is listed and their OM status summarized.

| Strain | Observations | | |
|---|---|---|---|
| | Constriction | PG fission (marked by IM fission) | OM fission |
| WT | 36 (OMs do not follow IMs in all 36 cell pairs) | 23 (20 with no significant OM invagination and 3 with significantly stretched OMs) | 2 |
| $Pal^{OE}$ | 26 (OMs follow IMs in all 26 cell pairs) | 12 (significant OM invagination in all 12 cell pairs) | 0 |
| $Lpp^{OE}$ | 19 (OMs follow IMs in all 19 cell pairs) | 14 (significant OM invagination in all 14 cell pairs) | 0 |
| $Lpp_{\Delta K}^{OE}$ | 18 (OMs do not follow IMs in all 18 cell pairs) | 15 (6 with no significant OM invagination and 9 with significantly stretched OMs) | 0 |

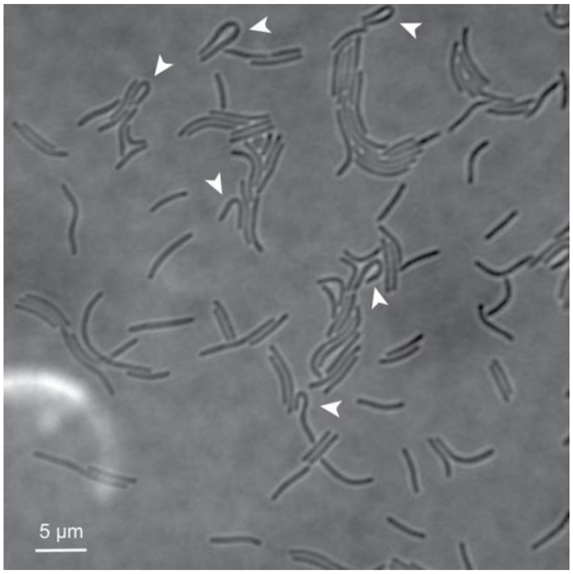


**Fig. S7. The flexibility of *M. xanthus* cell envelope.** Arrows point to the cells that bend significantly (>90°).

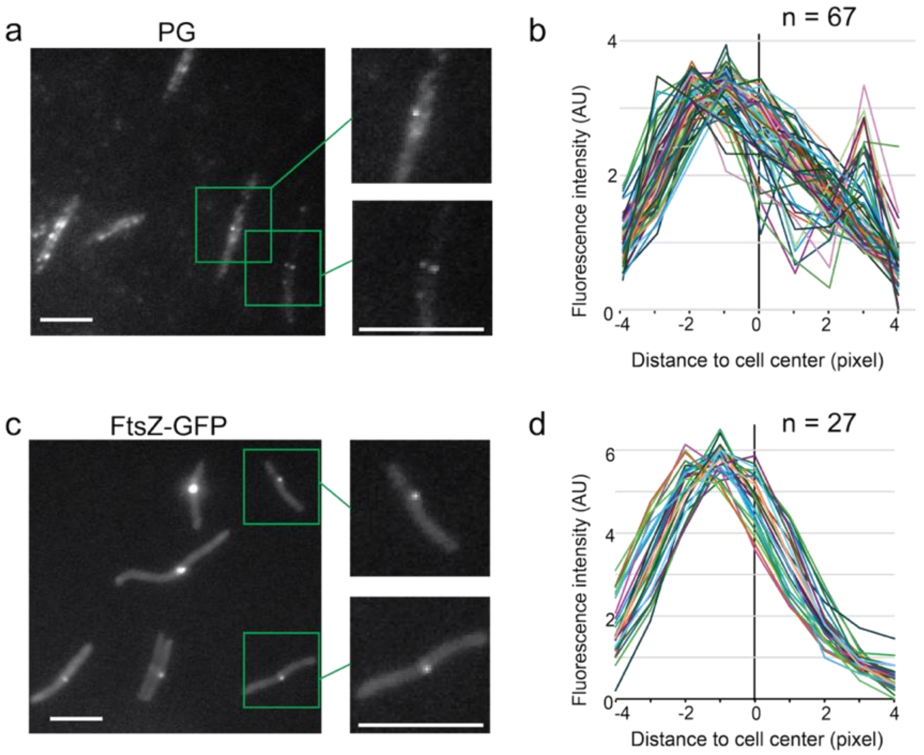


**Fig. S8. The asymmetric constriction of *M. xanthus* at division sites. a**) Asymmetric PG growth, as visualized by TADA, at the division sites of wild-type *M. xanthus*. **b**) The fluorescence intensities of TADA from 67 cells across cell width. **c**) FtsZ-GFP localizes in an asymmetric manner in dividing *M. xanthus* cells. **d**) The fluorescence intensities of FtsZ-GFP from 27 cells across cell width. In **b** and **d**, the centers of cells are designated as 0 and the identities of cells are distinguished by colors, and the fluorescence intensity values were normalized using Z-score standardization. Scale bars, 5 μm.

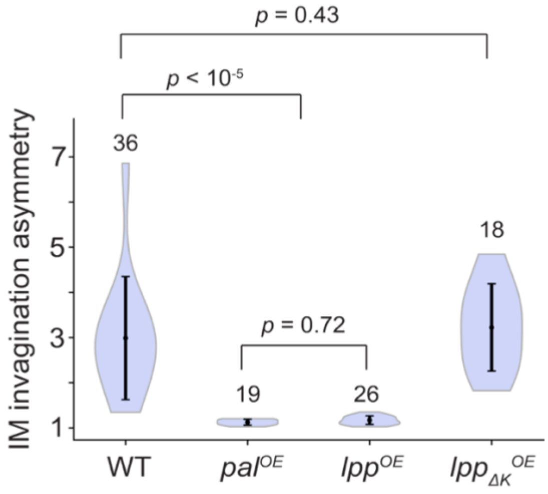


**Fig. S9. The asymmetry of IM invagination in the constriction stage.** Asymmetry is defined as the ratio between the IM-OM distances in cryoEM images at two sides of the division site. Dots and vertical bars are means and standard deviations, respectively. The number of cells measured is marked on the top of each plot and $p$ values are calculated using a one-way ANOVA test.

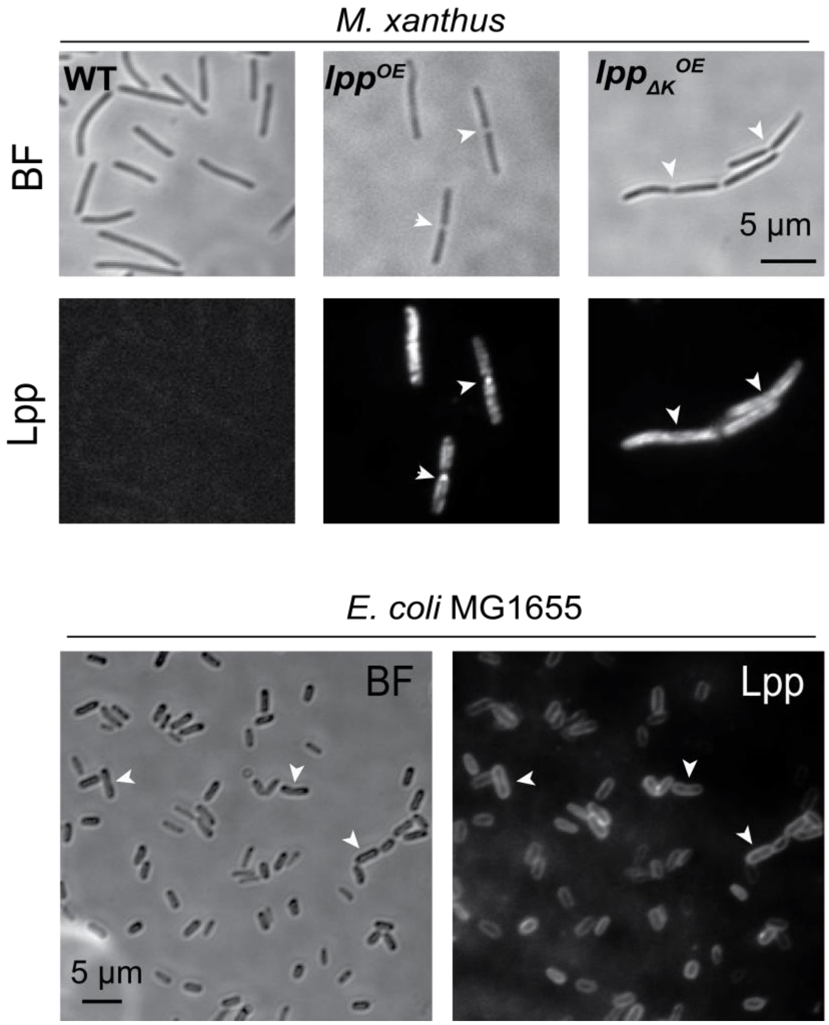


**Fig. S10. Exogenously expressed *E. coli* Lpp, but not the truncated Lpp$_{\Delta K}$, is enriched at the division sites of *M. xanthus*.** The localization of Lpp and Lpp$_{\Delta K}$ were detected by immunofluorescence using an anti-Lpp antibody. *E. coli* MG1655 that produces Lpp and wild-type *M. xanthus* that does not encode Lpp homologs were used as the positive and negative control, respectively, to show the specificity of the antibody. Cells were fixed and permeabilized on glass slides.

**Table S2. Strains and plasmids used in this study**

| Strain | Description | Source |
|---|---|---|
| DZ2 | The wild-type strain | [1] |
| DZ4469 | *pilA::tet* | [2] |
| BN333 | *ΔaglR pilA::tet* | [3] |
| BN380 | $P_{van}$-*lpp* | This study |
| BN381 | $P_{van}$-$lpp_{\Delta K}$ | This study |
| BN382 | $P_{van}$-*pal* | This study |
| BN383 | $P_{ftsZ}$-*ftsZ-gfp, Mx8 attB*, tet$^R$ | This study |
| BN384 | P*pilA-alkP-msfGFP Mx8 attB*, kan$^R$ | This study |
| **Plasmid** | | |
| pMR3629 | Plasmid carrying a vanillate-induced promoter, tet$^R$ | [4] |
| pMR3629-*lpp* | $P_{van}$-*lpp* in pMR3679, kan$^R$ | This study |
| pMR3629-$lpp_{\Delta K}$ | $P_{van}$-$lpp_{\Delta K}$ in pMR3679, kan$^R$ | This study |
| pMR3629-*pal* | $P_{van}$-*pal* in pMR3629, tet$^R$ | This study |
| pKA51 | $P_{ftsZ}$-*ftsZ-gfp, Mx8 attB*, tet$^R$ | [5] |
| pSWU19-*alkP-msfGFP* | pSWU19-P*pilA-alkP-msfGFP* kan$^R$ | This study |
| **Primers** | | |
| lpp-F | GGAATTCCATATGAAAGCTACTAAACTGG | This study |
| lpp-R | GGGGTACCCTAAGCAGCCTGAACGTCGGAACG | This study |
| pal-F | GGAATTCCATATGAACCGAACCCTCTTG | This study |
| pal-R | GGGGTACCCTACTCCCGCAGGCCCGTGG | This study |
| alkP-msfGFP_F | CGTAGTGCTAATCCCATCCGCGGGTCTAGAGGAAACCAAGAATAGAAAT AGAAAGGAGAATTAATGTTTCCTGCTGAGTGG | This study |
| alkP-msfGFP_R | CACCTTTACTGCCGCCGCCGCCGCTGGCGGAGGTGATGGTGCC | This study |
| msfGFP-Cmn_F | AGCGGCGGCGGCGGCAGT | This study |
| msfGFP-Cmn_R | GTCACGACGTTGTAAAACGACGCCAAGCTTTTATTTGTAGAGTTCATC | This study |

**Movies S1 – S6.** Time-lapse videos of dividing wild-type cells moving on a 1.5% agar surface. Images were captured at 10-s intervals and played at 10 frames/s (100 x speedup). Arrows point to the initial locations of the dividing cells. Scale bars, 5 μm.

**Movies S7, S8.** Time-lapse videos of dividing *pilA*$^-$ cells moving on a 1.5% agar surface. Images were captured at 10-s intervals and played at 10 frames/s (100 x speedup). Arrows point to the initial locations of the dividing cells. Scale bars, 5 μm.

**Movies S9, S10.** Time-lapse videos of dividing *ΔaglR pilA*$^-$ cells that are nonmotile on a 1.5% agar surface. Images were captured at 10-s intervals and played at 10 frames/s (100 x speedup). Scale bars, 5 μm.
